# Positional information encoded in the dynamic differences between neighbouring oscillators during vertebrate segmentation

**DOI:** 10.1101/286328

**Authors:** Marcelo Boareto, Tomas Tomka, Dagmar Iber

## Abstract

A central problem in developmental biology is to understand how cells interpret their positional information to give rise to spatial patterns, such as the process of periodic segmentation of the vertebrate embryo into somites. For decades, somite formation has been interpreted according to the clock-and-wavefront model. In this conceptual framework, molecular oscillators set the frequency of somite formation while the positional information is encoded in signaling gradients. Recent experiments using *ex vivo* explants have challenged this interpretation, suggesting that positional information is encoded in the properties of the oscillators, independent of long-range modulations such as signaling gradients. Here, we propose that positional information is encoded in the difference in the levels of neighboring oscillators. The differences gradually increase because both the amplitude and the period of the oscillators increase with time. When this difference exceeds a certain threshold, the segmentation program starts. Using this framework, we quantitatively fit experimental data from *in vivo* and *ex vivo* mouse segmentation, and propose mechanisms of somite scaling. Our results suggest a novel mechanism of spatial pattern formation based on the local interactions between dynamic molecular oscillators.

## Introduction

Pattern formation during embryonic development requires that the cells assess their spatial position. One useful conceptual framework to understand this process is to assume that each cell has a positional value that relates to its position in the coordinate system. The cells then use this positional information to coordinate their differentiation process (Wolpert, 1969). Based on this conceptual framework, Cooke and Zeeman proposed a model to explain the sequential and periodic formation of the somites in the vertebrate embryo (Fig. 1A). In their model, the positional information of the cells is given by a signaling wavefront, while a clock sets the frequency of somite formation (Fig. 1B) (Cooke and Zeeman, 1976).

**Figure 1.**
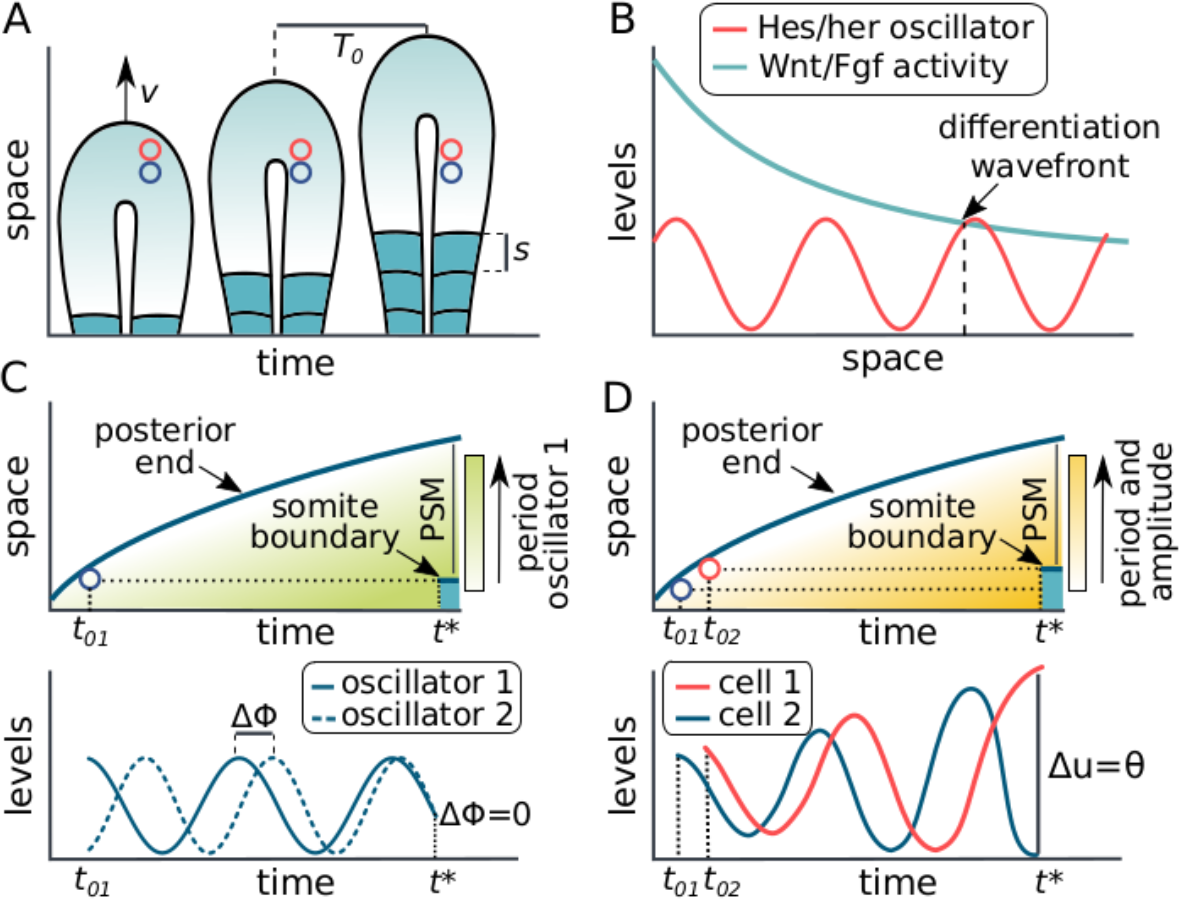
Graphic representation and models of vertebrate segmentation. A) During embryonic development, the tail of the embryo extends due to the incorporation of new PSM cells (red and blue circles) with a velocity (v), and new segments (s) are formed periodically (period=T_0_). B) Representation of the clock-and-wavefront model: a positional posterior-anterior differentiation front (or wavefront) is created by a gradient of Wnt and Fgf activity. As the tail growths, the cells cross the wavefront and are incorporated into a new somite. While the wavefront defines the position of the new somite (dotted vertical lines), the oscillatory expression of Notch genes forms a clock that defines the period of segmentation. C) Representation of the phase-difference model: each cell has two oscillators, one oscillator with a dynamic period (green area) and one with a fixed period. Positional information is encoded in the differences in phase between the oscillators in each cell. Somite formation will occur when the shift in the phase becomes sufficiently small, as represented at t=t*. D) Representation of the level difference model: positional information is encoded in the differences in the levels of neighboring oscillators. Neighboring cells are added to tissue at different time points (t_0_) and start with slightly different levels of the oscillator. These levels accumulate due to differences in the period and amplitude (yellow area). Somite formation will occur when the difference in the levels (Δu) exceeds a threshold (θ), as represented at t=t*.

The clock-and-wavefront model guided much of the experimental work done in the subsequent decades. The rhythm of segmentation has been shown to be accompanied by travelling waves of gene expression, which sweep from the tail bud to the anterior end of the presomitic mesoderm (PSM) (Masamizu et al, 2006). These waves emerge from the oscillatory expression of ‘clock’ genes involved in the Notch pathway, such as *Lfng* and *Hes* (McGrew et al 1998; Palmeirim et al., 1997; Forsberg et al, 1998; Dequeant et al., 2006; Niwa et al., 2007). In the tail bud region, where the PSM cells are generated, *Fgf8* and *Wnt3* are produced and as the cells cross the PSM their mRNA levels decrease due to degradation, creating a gradient by inheritance (Dubrulle and Pourquié 2004; Aulehla et al, 2003). Perturbations on Wnt and Fgf gradients have been shown to affect somite formation (Dubrulle et al, 2001; Sawada et al, 2001; Naiche et al 2011), as required if the signaling gradients encode the wavefront of somite formation.

Recently, experiments of *ex vivo* explants have challenged the clock-and-wavefront view and favored an interpretation where the wavefront is implicit in the dynamics of the clock (Lauschke et al, 2013; Tsiairis and Aulehla 2016). In these experiments, tail bud tissue is explanted and forms a monolayer PSM (mPSM) with concentric travelling waves sweeping from the center of a dish to its periphery. Interestingly, after growth stops, segments begin to form, and the size of the segments scales with the size of the mPSM (Lauschke et al 2013). The formation of these segments in the absence of growth cannot be easily explained with the clock-and-wavefront model as it would require long-range mechanisms that sense the mPSM length.

Alternative models of somite formation have been proposed, as recently reviewed (Pais-de-Azevedo et al 2018). Among these models, the phase-difference model is able to explain segment scaling as observed in *ex vivo* explants. In this model, positional information is encoded in the phase of the oscillator (Goodwin and Cohen, 1969). Experiments using *ex vivo* explants have shown that the difference in phase between the cells in the center of the explant and the cells in the newly formed segment is constant and equal to 2*π*, supporting the idea that the phase of the oscillator alone is a predictive parameter for the position of somite formation (Lauschke et al., 2013). However, this constant phase difference is not observed *in vivo*, as measured in zebrafish (Soroldoni et al, 2014). Therefore, for this model to work *in vivo*, an additional oscillator would be required in each cell and the wavefront would then be defined via the relative phase between these two cellular oscillators (Figure 1C, Lauschke et al., 2013). Network architectures that can compute such a relative phase of oscillation have been investigated (Beaupeux and François, 2016), and recent experiments show that the phase shift between Notch and Wnt signaling can control segmentation in *ex vivo* explants (Sonnen et al, 2018). It remains to be shown whether this mechanism can also explain *in vivo* segmentation.

The *Hes/her* genes, which are key targets of the Notch signaling pathway (Takke and Campos-Ortega, 1999; Bessho et al, 2001; Kageyama et al, 2007), are the central components of the cellular oscillators that control somite formation. In various species, these genes have been shown to oscillate due to a delayed autoinhibition, although there is a substantial variability in the gene network (Krol et al, 2011; Hirata et al 2002; Oates and Ho 2002; Lewis 2003). In many species, the period of Hes/her oscillations increases in time as the cells move from the posterior to the anterior part of the PSM (Delaune et al, 2012; Shih et al, 2015; Gomez et al, 2008; Tsiaris and Aulehla, 2016). The same is true for the amplitude of the oscillators, as quantified in zebrafish (Delaune et al, 2012; Shih et al, 2015), and indirectly shown in mouse (Lauschke et al, 2013; Tsiaris and Aulehla, 2016). Interestingly, the temporal increase in period and amplitude correlates with the temporal decrease of Fgf and Wnt signaling due to mRNA degradation. Wnt activity has indeed been found to modulate the period of Hes/her oscillations in PSM cells (Gibb et al, 2009; Wiedermann et al, 2015; Dubrulle et al, 2001; Sawada et al, 2001), but it remains to be tested whether Wnt activity also modulates the amplitude of these oscillators.

Here, we show that the positional information during somite formation can be encoded by a single molecular oscillator if its period and amplitude increase in time and space as measured for the *Hes/her* oscillator. A set of neighboring oscillators whose period and amplitude follow such a gradient generate travelling waves. This can be visualized in a pendulum wave experiment, where a set of pendulums with gradually increased lengths that start from the same initial condition display a travelling wave (Berg, 1991; Flaten and Parendo, 2001). We now show that the different levels of neighbouring oscillators can encode positional information during somite formation (Fig. 1D). Here, only a single molecular oscillator (Hes/her) is required per cell. When the PSM cells are incorporated in the tail bud region, they start with the same initial condition and Hes/her oscillations are synchronized with their neighbors. As the cells leave the tail bud region and cross the PSM, a temporal increase in the period and amplitude of Hes/her oscillations leads to a gradual increase in the difference of Hes/her levels in neighboring cells. Somite boundaries can then be triggered by a critical difference of Hes/her levels between neighbouring cells (Figure 1D). In the following, we first develop a theoretical framework for the proposed mechanism. We then show that our model quantitatively fits *in vivo* mouse segmentation from wild type and growth-perturbed embryos, captures temperature compensation in somite size and predicts a delayed scaling between somite size and PSM length. We further use data from different species to validate the predicted scaling between somite size and PSM length, and to suggest possible developmental mechanisms of somite size control. Lastly, we quantitatively fit data from mouse *ex vivo* explants, showing that the proposed mechanism can in principle explain data from both *in vivo* and *ex vivo* mouse segmentation.

## Results

### Theoretical framework

In our framework, each cell expresses an oscillatory protein *u* that represents a member of the Hes/her family. For simplicity, we represent the levels of *u* by a sine function:

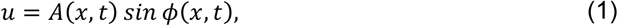

where the amplitude *A* and the phase ϕ depend on the position of the cell *x* and the time *t*.

After leaving the tail bud region, the amplitude and period of the oscillations increase over time in the PSM cells (Delaune et al, 2012; Shih et al, 2015). The exact functional form of this increase has not yet been determined. For convenience, we mathematically represent the increase in the amplitude (A) and period (T) by exponential functions:

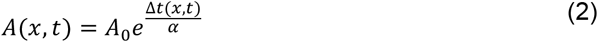

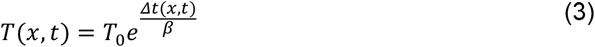

where *A*_*0*_ and *T*_*0*_ are the amplitude and period at the tail bud region, respectively, and α and β are the characteristic time scale of amplitude and period gradients, respectively. Note that the period and, consequently, the frequency are dependent on the amount of time a cell has spent in the tissue. This time interval is given by Δ*t(x,t)* = *t* − *t*_*0*_, where *t*_*0*_ = *t*_*0*_(x) represents the moment the cell in position *x* is incorporated into the PSM (Figure 2A). The body axis elongates mainly by growth at the tail bud. Accordingly, the tail bud moves in the direction of increasing *x*, while the position of each cell, *x*, in the PSM can be considered as fixed (Figure 2A).

**Figure 2.**
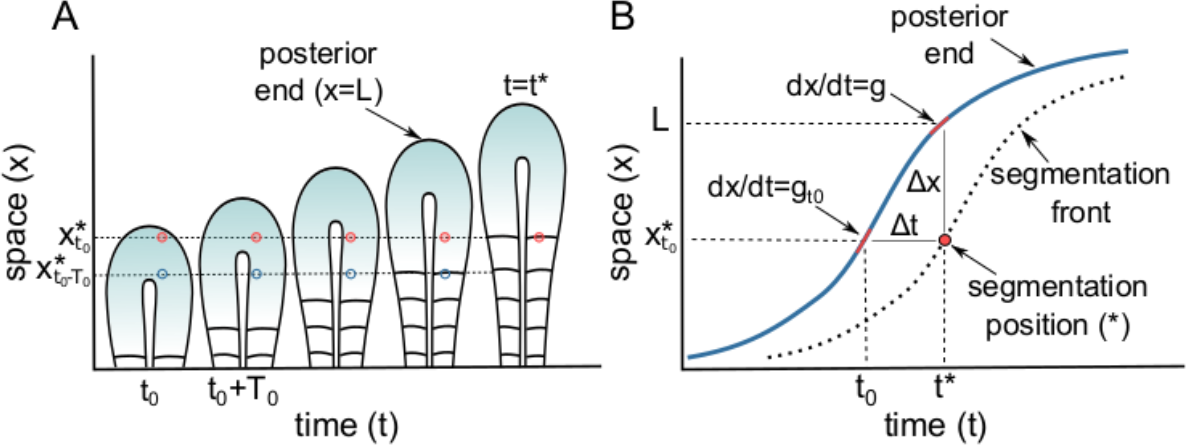
Schematic representation of segmentation process. A) The tail of the animal grows as new cells are added into the tail bud. Red dot represent a cell that is added into the tail bud at time (t=t_0_) and incorporated into the PSM at time (t=t*). B) The blue curve represents the position of the posterior end in time, the red dot represents the position of segmentation where du/dx=θ. The variable Δt=t−t_0_ represents the amount of time since the cell at position x* was incorporated to the tissue and Δx=L-x* is the distance of the cell x* to the posterior end at the time t. The tail bud growth rates g_t0_ and g represent the growth rate at the tail bud at the time t_0_ and t*, respectively. Note that t_0_ = t_0_(x) such that 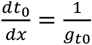.

The phase *ϕ* of the cellular oscillator is related to the period (*T*) and the frequency (*ω*) of the oscillation according to

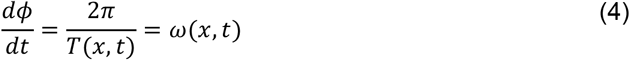

which can be integrated to yield

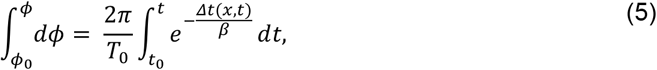

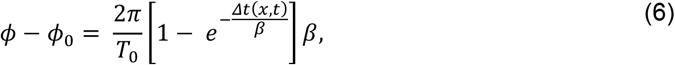

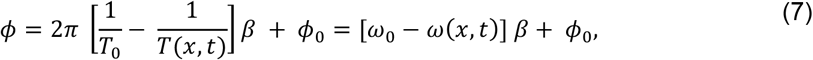

where *ϕ*_0_ = *ω*_*0*_*t*_*0*_*(x)* represents the initial phase and 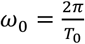 the initial frequency of the oscillator at the tail bud.

The slight difference in period and amplitude between neighboring cells leads to the temporal accumulation of differences in their levels *u*. We propose that segmentation occurs when these differences in *u* reach a certain threshold (*θ*). This can be represented mathematically as:

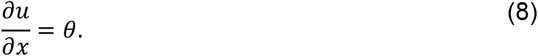

The derivative of *u* can be written as

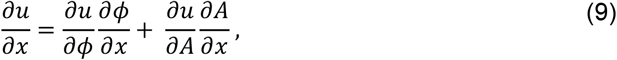

which leads to

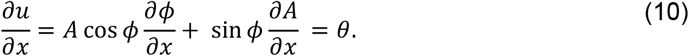

By calculating the spatial derivatives 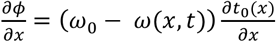 and 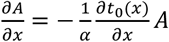, we have:

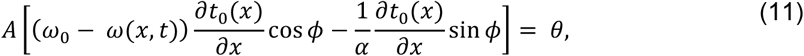

where the derivative

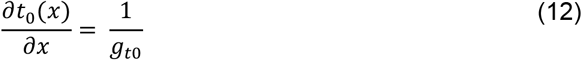

is the inverse of the tail bud growth rate, *g*_*t*0_, at time *t*=*t*_*0*_ (Figure 2B). Combining Eqs. 11 and 12, we obtain:

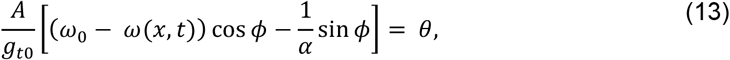

which can be rewritten as:

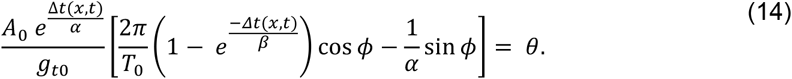

This implicit equation yields the time interval, *Δt* = *t* − *t*_*0*_*(x)*, between the time a cell leaves the tail bud and the time it forms a segment (Fig. 2). As the tail bud is extending posteriorly with growth rate *g*, the time interval *Δt* determines the distance *Δx* between the posterior end *L* and the segment boundary *x* (PSM length), as well as the distance between the previous and the new somite boundary, i.e. the size of the newly formed somite (Figure 2). Importantly, when forming a new somite, posterior cells reach the threshold before the anterior cells and the anterior part of the new somite experience high levels of the oscillatory protein (Figure S1), consistent with experimental observations in zebrafish (Shih et al, 2015).

The model has only 5 parameters: the characteristic time scale of the amplitude gradient (α), the characteristic time scale of the period gradient (β), the tail bud growth rate (*g_t0_)*, the oscillation period at the tail bud (T_0_), and the normalized threshold for segmentation (θ/A_0_). Importantly, these parameter values are all set at the time *t*_*0*_ when the cells leave the tail bud region. Consequently, after the cells have left the tail bud, only *Δt* changes. This leads to a timer mechanism of somite formation, independent of any spatial input.

### Model validation with quantitative data from growth-perturbed mouse somitogenesis

During mouse somitogenesis, the axial growth rate changes substantially and follows a hump-shaped curve, an initial increase followed by a consecutive decrease (Tam 1981, Figure S2). As a consequence of such growth profile, the axial length increases in a sigmoid-like fashion during development (Tam 1981, Figure S2). In addition to the axial growth rate, also the somite sizes and the PSM length change substantially during embryonic development and are disturbed in animals that suffered drastic size reduction due to treatment with DNA-synthesis inhibitor Mitomycin C (MMC) (Tam 1981). Interestingly, although growth-perturbed (MMC-treated) embryos are significantly smaller than wild type (WT) at early stages, these embryos show compensatory growth, resulting in an embryo with normal final size (Tam 1981, Figure S2). Such compensatory growth, however, leads to a disturbed somitogenesis where both the PSM length and somite size are smaller compared to WT embryos (Tam 1981, Figure S3). We sought to use this data to test whether our model would be able to correctly reproduce the measured changes in somite sizes and PSM lengths for the different growth rates at the different embryonic stages, and in the different growth conditions.

The time-dependent tail bud growth rate *g(t)* can be inferred from the experimental data and is then used as input to the model (Tam 1981, Figures 3A and S2). When we keep all other parameters fixed during the segmentation process, the model fails to reproduce somite size and PSM length (Tam 1981, Figure S4 A-C). In the next step, we investigated the case where the amplitude and period in the tail bud is not constant over time, but depends on the varying growth rate *g*_*t0*_ in the tail bud (Figure 3A). We use an exponential relationship

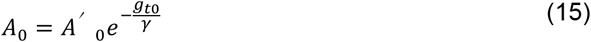

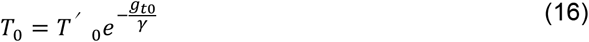

with an additional parameter γ. Eq. 14 remains valid since both *g*_*t0*_ and γ are independent on *x*. The extended model fits the segmentation period (Figure S4) as well as the measured PSM lengths (Figure 3C,D; Fig. S4E) and somite sizes (Figure 3E,F; Fig. S4F), both in control and MMC-treated embryos. The difference in the growth rate between WT and MMC-treated embryos (Figure 3A,B) alone is, however, not sufficient to explain the differences in somite size and PSM length between these two conditions (Figure 3G). In particular, the characteristic time scale of the amplitude gradient (*α*) must additionally differ between these conditions.

**Figure 3.**
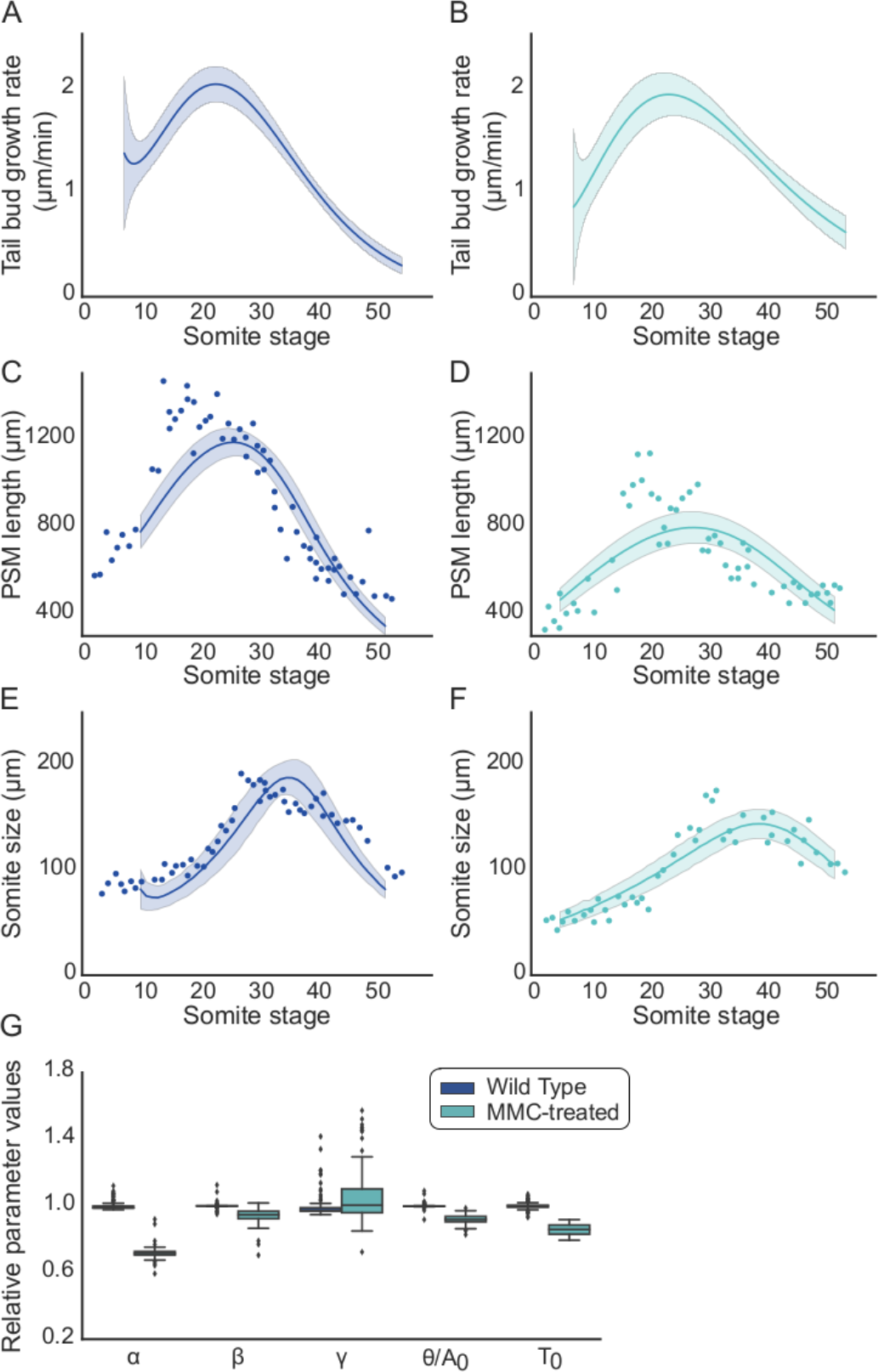
Model validation with segmentation data from control and MMC-treated mouse embryos. A) Inferred tail bud growth rate, g, for different somite stages of WT and B) MMC-treated embryos. Tail bud growth rates were inferred from experimental data (Tam 1981, Figure S2). Lines represent the average fit of a bootstrap resampling and the area represents the 95% confidence interval. C) PSM length for different somite stages for WT and D) MMC-treated embryos. E) Somite size for different somite stages for WT and F) MMC-treated embryos. C-F) Dots represent data from (Tam 1981), lines represent the average fit and the area represents the 95% confidence interval of the model prediction. A total of 100 simulations were evaluated using different fits of the tail bud growth rate, obtained via bootstrap, as an input. G) Comparison of parameter values that best fit the data for WT and MMC-treated embryos. Values are normalized by the average values of WT embryos, which are presented in Table 1.

These results show that our framework quantitatively reproduces mouse segmentation *in vivo* as long as a modulation in the growth rate additionally affects the characteristic time scale of the amplitude gradient, suggesting a link between the growth rate and the properties of the cellular oscillators.

### Somite size and PSM length are determined at the tail bud and depend on the time-varying growth rate

Somite size scales with body size (Cooke 1975) and even in *ex vivo* explants the segment size scales with mPSM size (Lauschke et al, 2013). In addition, measurements in mouse reveal an intriguing temporal relationship between the growth rate, PSM length and somite size: all three curves follow a hump shape (Tam 1981, Figure 3). The peak of the growth rate coincides approximately with the peak of PSM length, while the peak of the somite size is delayed in relation to growth rate and PSM length (Tam 1981, Figures 3, S2,3). What determines the relative size of somites and PSM? And what determines this delay?

The size of a somite (s) is defined by the difference between the position of the new segment formed at t=t* (x*_t_) and the position of the new segment formed at t=t*-T_0_ (x*_t-T0_). The position of segment formation can be estimated by integrating the growth rate until the moment the cell is incorporated into the tissue t=t_0_ (Figure 2A),

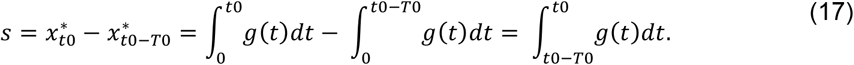

Assuming that the growth rate is constant in the time period [t_0_−T_0_, t_0_] that corresponds to one somite stage, the size of the somite is approximately

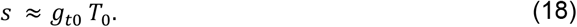

The size of the somite, *s(t)*, that is formed at time *t* thus depends on the growth rate, *g*_*t0*_, and the period, *T*_*0*_, when the cells are incorporated into the PSM.

What about the PSM? According to our model, the size of the PSM (*P*) is directly proportional to the tail bud growth rates

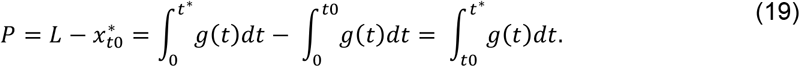

Assuming again that the growth rate is constant in the time period [t_0_, t*], the size of the PSM is approximately:

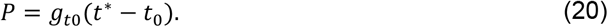

Accordingly, the PSM length achieves its maximum soon after the growth rates peaks, which is consistent with the mouse data (Tam 1981; Figure 3).

So how long is the delay between the maximal somite size and the maximal growth rate? This delay is proportional to the time it takes the cells from being added into the PSM (t=t_0_) to be incorporated into a new somite (t=t*). One simple way to estimate this is to count how many somites have to be formed until the cells that are in the tail bud become incorporated into a new somite. This can be estimated by dividing the size of the PSM by the size of the somites, giving a good estimate in somite stages (Figure 2A). Another way to estimate this delay is to use equations 18 and 20 as a constant growth rate is a good approximation around the somite stages where the growth rate is maximal. In this case, we obtain that the delay (*τ*) is given by:

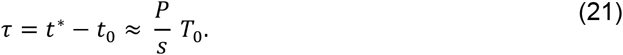

Therefore, our model predicts a delayed scaled relationship between the PSM length and somite size, where the delay is approximately the ratio PSM length to somite size (P/s). To confirm that this relationships indeed holds when growth rates change over time, we considered an idealized hump-shaped tail bud growth rate curve (Figure 4A): the difference in the peaks of PSM length and somite size approximately corresponds to the amount of time the cells spend to cross the PSM, which is approximately the ratio *P/s* (Figure 4B). Finally, we compared the measured differences in the peaks of somite size and PSM length for mouse, chicken and snakes: as predicted by our model, this difference, in somite stages, is approximately the ratio *P/s* (Figure 4C, Gomez et al, 2008).

**Figure 4.**
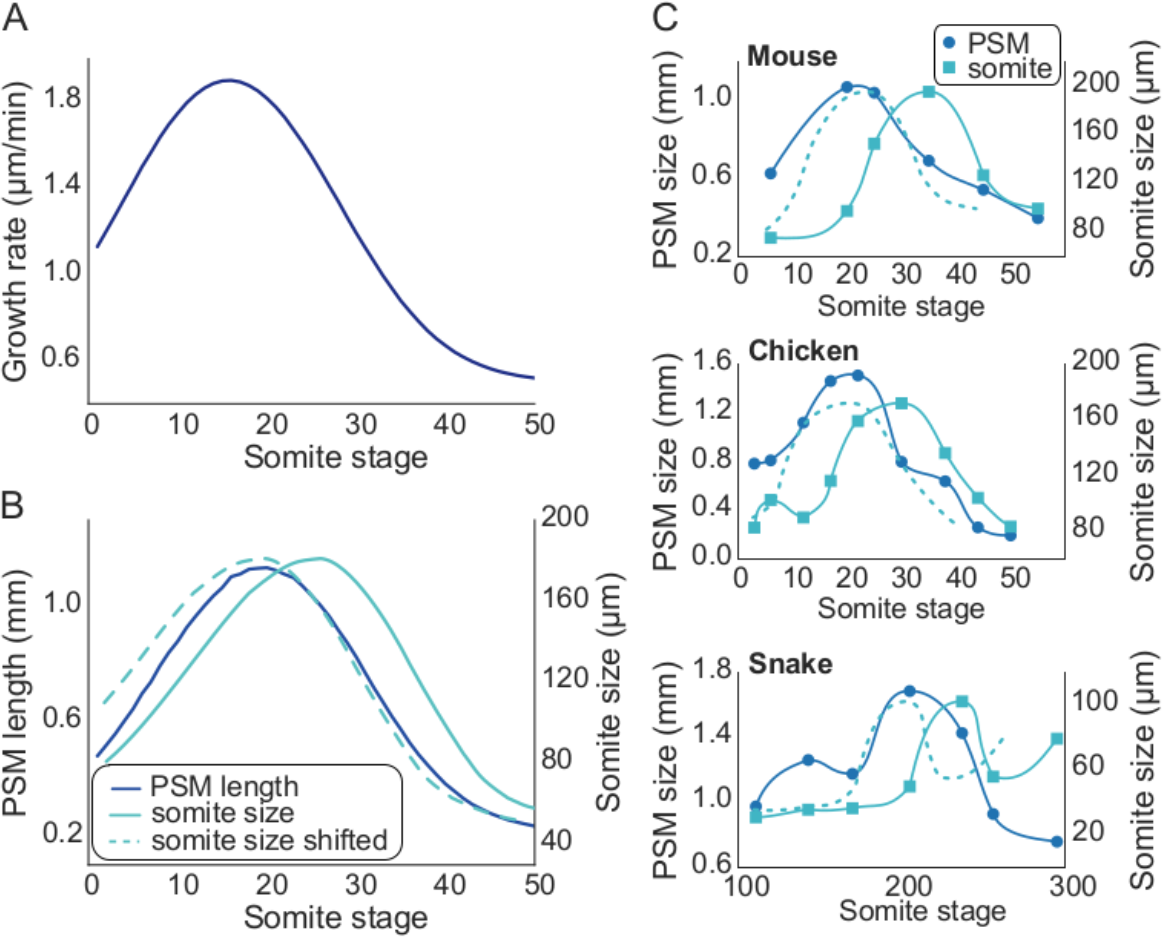
Relationship between tail bud growth rate, somite size and PSM length for dynamic growth rates. A) Tail bud growth rate for different somite stages. B) Inferred PSM length (blue curve) and somite size (green curve) for the tail bud growth rate presented in A) and with parameters consistent with mouse segmentation (Table 1). The dashed green line represents the somite size curve shifted by the ratio PSM/somite size at the somite stage when PSM length is maximum. C) Experimental measurements of PSM length and somite size during different embryonic stages for mouse, chicken and snake obtained from (Gomez et al, 2008). The dashed curve represents the somite size curve shifted by the ratio PSM/somite size at the somite stage when PSM length is maximum.

These results support the idea that the size of the somites depends on the growth rate at the time the cells are at the tail bud. Such a relationship is consistent with our timer-based model (Eq. 14) where all properties are defined at the moment when the cells enter the PSM and somite formation does not require additional spatial inputs, but would not be consistent with the clock-and-wavefront model with its positionally controlled segmentation front. The clock-and-wavefront model reproduces Eq. 18 only in case of a constant growth rate, while our model can explain the observed relationship between PSM length and somite size also for time-varying growth rates. A timer mechanism, as we propose, is also consistent with observations in zebrafish where the determination front is defined already a few somite stages before segmentation occurs (Akiyama et al, 2014). Similarly, PSM cells in the mouse are organized into segmental units before segmentation (Tam 1981).

### Evolutionary mechanisms of vertebrate segmentation

There is a large diversity in somite size, number and frequency among vertebrate species, but little is known about the evolutionary mechanisms that lead to such diversity (Gomez et al, 2009). For example, while mouse and chicken share similar segmentation properties, snakes and zebrafish have somites three to four times smaller (Figure 5A). Interestingly, snakes and zebrafish achieve smaller somites via different mechanisms. In zebrafish, smaller somites are mostly due to a decrease in the period of oscillations, while snakes have smaller somites due to slower growth rates (Gomez et al, 2008; Figure 5A). This suggests that differences in the growth rate and period of oscillations are the main evolutionary mechanism to achieve somites with different sizes. But are the changes in these parameters sufficient to explain segmentation properties of these different species? To answer that, we used our model to estimate somite size and PSM length for different values of growth rate and oscillation period in the tail bud. We found that changes in these parameters alone are sufficient to explain somite size, but not the observed PSM length (Figure S5).

**Figure 5.**
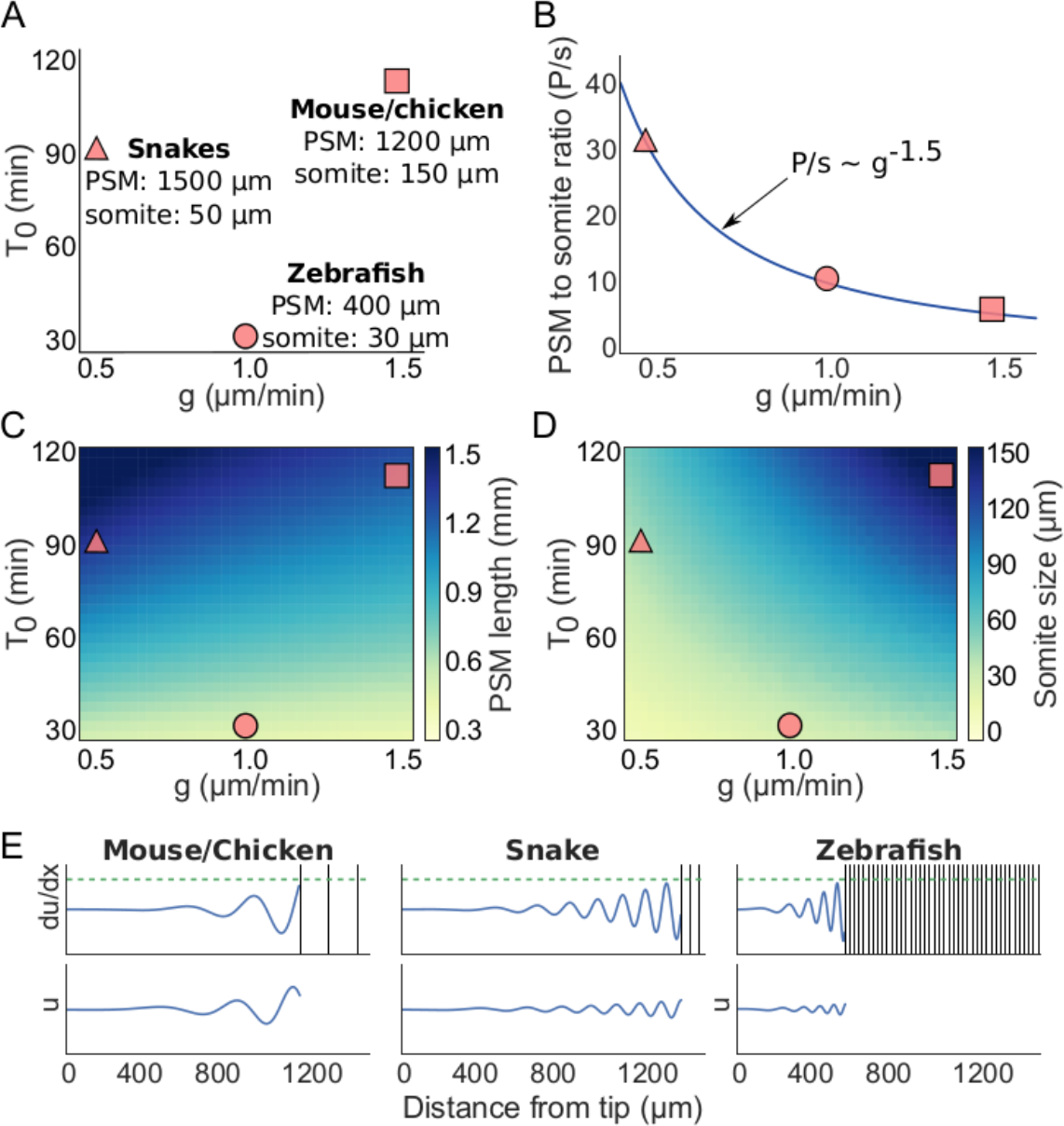
Segmentation properties of different vertebrate species. A) Representation of the estimated overall growth rate and period in the tail bud, PSM length and somite size for mouse and chicken, snakes and zebrafish (Gomez et al, 2008). B) Relationship between PSM/somite size ratio (P/s) and the tail bud growth rate for different species. Blue line represents the fit that relates the ratio P/s to the growth rate. C) PSM length and D) somite size for different values of growth rate and period at the tail bud. We considered that changes in the growth rate and the period of oscillations also affect the steepness of the gradients. E) Representation of the values of u and ∂u/∂x for different species. Dashed green lines represent the threshold θ, while dark vertical lines represent the position of formed segments.

We further asked if concomitant changes in the characteristic length of the gradients (α and β) could explain changes in both somite size and PSM length. The differences in the period at the posterior end and at the anterior part of the PSM scales with PSM length in a similar fashion between snakes and zebrafish (Gomez et al, 2008). This suggests that the slowing down of the oscillations is similar in different species:

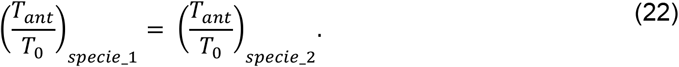

Consequently:

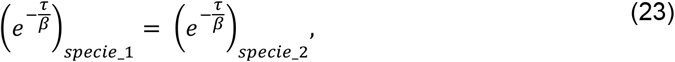

which leads to a relationship between the characteristic time scale of the period gradient (β) and the time the cells take to cross the PSM (т):

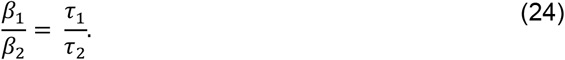

As previously discussed (Eq. 21), the time the cells take to cross the PSM can be inferred in terms of the PSM length and somite size:

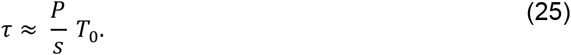

Using data from different species, we inferred the following relationship between *P/s* and *g* (Figure 5B):

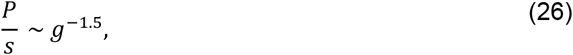

and by combining Eqs 24–26, we can then estimate the inter-species ratio of the characteristic length of the period as a function of changes in the growth rate and period at the tail bud:

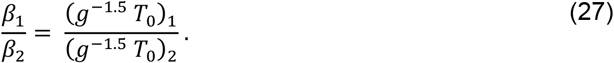

Assuming that the inter-species ratio of the characteristic length of the amplitude follows the same relationship, we have:

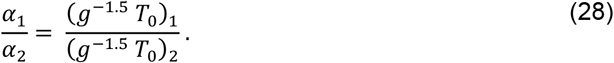

If we derive the α and β for the other species with equations 27 and 28 based on the α and β that we previously inferred from experimental data for the mouse (Figure 3, Table 1), we correctly estimate the somite size and PSM length for mouse, chicken, snake and zebrafish (Figure 5C-D). In addition, we observe a much larger number of stripes of the oscillatory protein in the PSM of snakes compared to zebrafish, mouse and chicken, which is consistent with experimental observations (Figure 5E; Gomez et al, 2009).

These results suggest a developmental mechanism of somite size control, where a decrease in the growth rate leads to an increase in the characteristic length of the amplitude and period gradients (α and β), which consequently leads to an increase PSM length to somite size ratio (P/s). We predict that the ratio P/s increases as the tail bud growth rate decreases (Eq. 26). It would be interesting to measure these parameters in other species to further confirm the existence of such a relationship.

### Three model parameters mainly determine somite size, PSM length, and the segmentation period

In our analysis above, we noticed differences in how the growth rate, *g*, and the characteristic length of the amplitude gradient (*α*) affected somite size and PSM length. To discern the individual impact of each parameter in our model, we carried out a parameter sensitivity analysis. To this end, we performed a perturbation on the values of each parameter and assessed the effect on somite size, PSM length, and the segmentation period. Here, we used a constant tail bud growth rate, *g*, such that the PSM length and somite size do not change during the segmentation process (Eq.17–20) (Figure 6A,B). To study the individual effects of parameters, we used the model where the period and amplitude are independent of the tail bud growth rate (Eq. 2,3), but the same conclusions hold also when they are dependent (Eq. 15,16) (Figure S6). The sensitivity analysis reveals that PSM length, somite size and segmentation period are controlled mostly by three key parameters: the tail bud growth rate (*g*), the oscillation period at the tail bud (*T*_*0*_), and the characteristic length of the amplitude gradient (*α*). Here, *g* positively regulates both PSM length and somite size, *T*_*0*_ positively regulates both the somite size and segmentation period, and *α* positively regulates the PSM length (Figure 6C).

**Figure 6.**
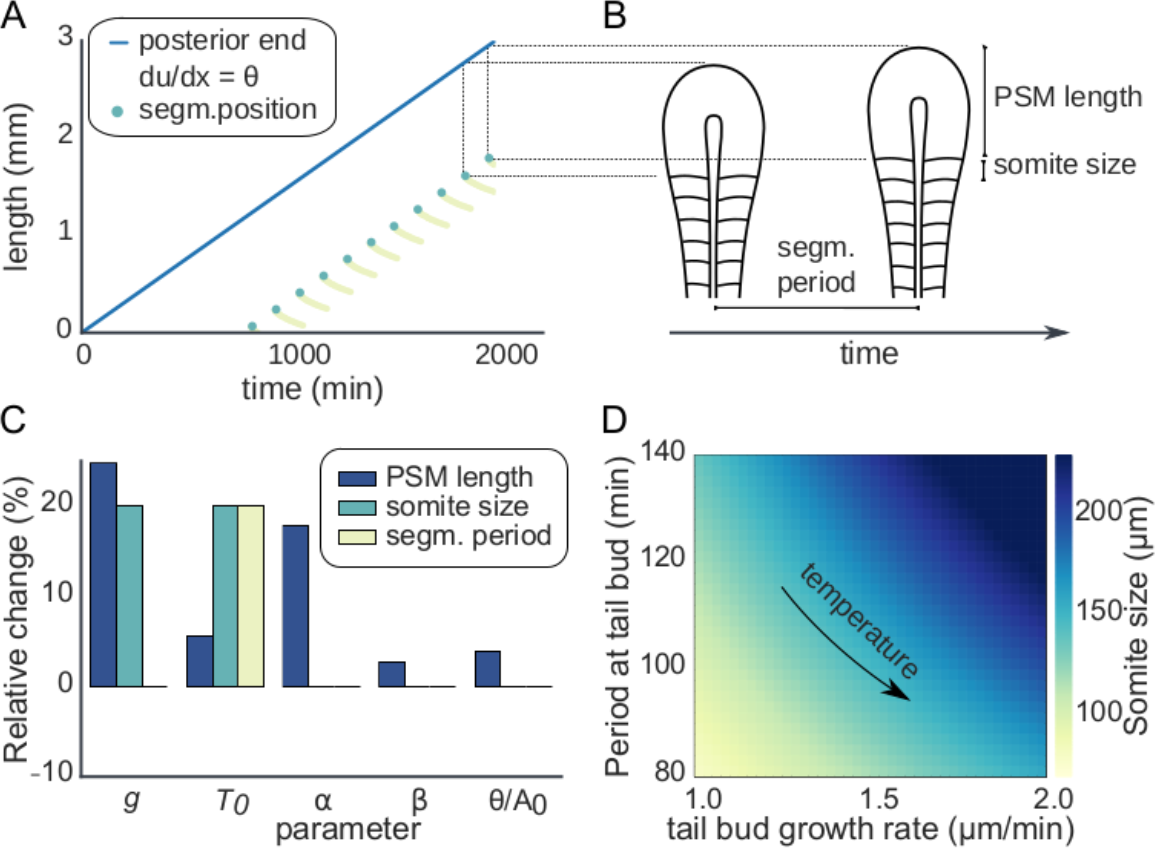
Segmentation properties at constant tail bud growth rate. A) Position of posterior end (blue line) and position of segmentation in time. Yellow dots represent the points where ∂u/∂x = θ and green dots represent where a new segment is formed. The formation of a new segment happens when ∂u/∂x = θ is satisfied in a position x such that x > x_s_, where x_s_ is the position of the previous segment. The phase of the cells during segmentation is the same during the whole process (Figure S1). Note that caudal cells reach the threshold before the rostral cells within a somite length. B) Graphical representation of segmentation process. Assuming constant tail bud growth rate, the PSM lengthening is exactly the size of one somite during one segmentation period. C) Parameter sensitive analysis. Each parameter is increased in 20% of its standard value (Table 1). The tail bud growth rate (g), clock period at posterior end (T_0_), and the characteristic length of amplitude gradient (α) are the most sensitive parameters, i.e., lead to changes of more than 10% of either PSM length, somite size and segmentation period. PSM length is regulated by both g and α, somite size is regulated by both g and T_0_, and segmentation period is controlled by T_0_. D) Somite size for different values of T_0_ and g. If an increase in g is accompanied by a proportional decrease in T_0_, the size of somites remains the same, suggesting a temperature compensation mechanism.

### Somite size is temperature compensated and controlled by tail bud growth rate and clock period

Intriguingly, somite size in zebrafish remains the same when embryos are grown at different temperatures, even though the growth rates and oscillation periods change substantially with temperature (Oates et al, 2012). In fact, according to the clock-and-wavefront model and consistent with experimental data, the combined changes in the growth rate and the oscillation period compensate in a way that the somite size remains constant (Schröter et al, 2008; Oates et al, 2012). Also, in our model, the size of the somite is affected in the same way by the tail bud growth rate and the oscillation period (Figure 6C), and the somite size remains constant when the growth rate and the oscillation period in the tail bud are changed in parallel (Figure 6D). This further suggests that a molecular mechanism exists that couples the oscillation period and probably also the oscillation amplitude to the growth rate (Eq. 15,16).

### Pattern scaling of *ex vivo* explants with different temperatures

Cells from the PSM self-organize into monolayer PSM (mPSM) structures when explanted *in vitro* (Lauschke et al, 2013; Tsiairis and Aulehla 2016). These structures show many properties that resemble *in vivo* segmentation. For example, the activity of Wnt and Fgf is higher in the cells in center of the tissue and lower in the cells close to the periphery, forming a center to periphery gradient which is similar to the posterior to anterior gradient observed *in vivo*. Also, a period and amplitude gradient from center to periphery is observed. Moreover, segmentation is observed from the periphery to the center of the tissue, leading to a gradual shrinkage of the mPSM. Interestingly, the size of the segments scales with the size of the remaining mPSM. In contrast to what is observed *in vivo*, however, there is no growth during *ex vivo* segmentation and the cells stay in a fixed position in relation to the center of the mPSM.

While the period gradient in mPSM explants has previously been described by a time-dependent function (Lauschke et al, 2013), we noted that the data can as well be described with a period gradient that only varies in space, but not in time (Supplementary Information, Figures S8,9). In the spirit of parsimony, we therefore assume that the period and amplitude gradients change only over space, but not with time; we note that a time-varying period gradient would yield similar results. We thus describe the amplitude and period gradients by

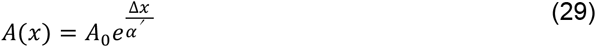

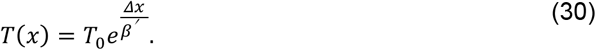

Here, α’ and β’ are the characteristic spatial length of the amplitude and period gradients, respectively, and *Δx* = *L* − *x*, is the distance of the cell x to the center of the mPSM (*L*). The position of segmentation is then be determined by:

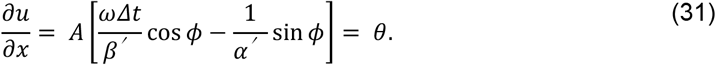

We emphasize that in case of a static period gradient, the phase of the oscillators, *ϕ*(*x*) = *ω*(*x*) Δ*t* changes in time. We then use Eq. 31 to fit the relationship between somite size and wave velocity with PSM length for explants at different temperatures (Figure 7A-E; Lauschke et al, 2013). We noted that most parameters do not change, except the period of oscillations, T_0_, which must be longer for explants at lower temperature (Figure 7E). This is consistent with experimental measurements showing that explants at lower temperature have a longer overall oscillation period (Figure 7F; Lauschke et al, 2013).

**Figure 7.**
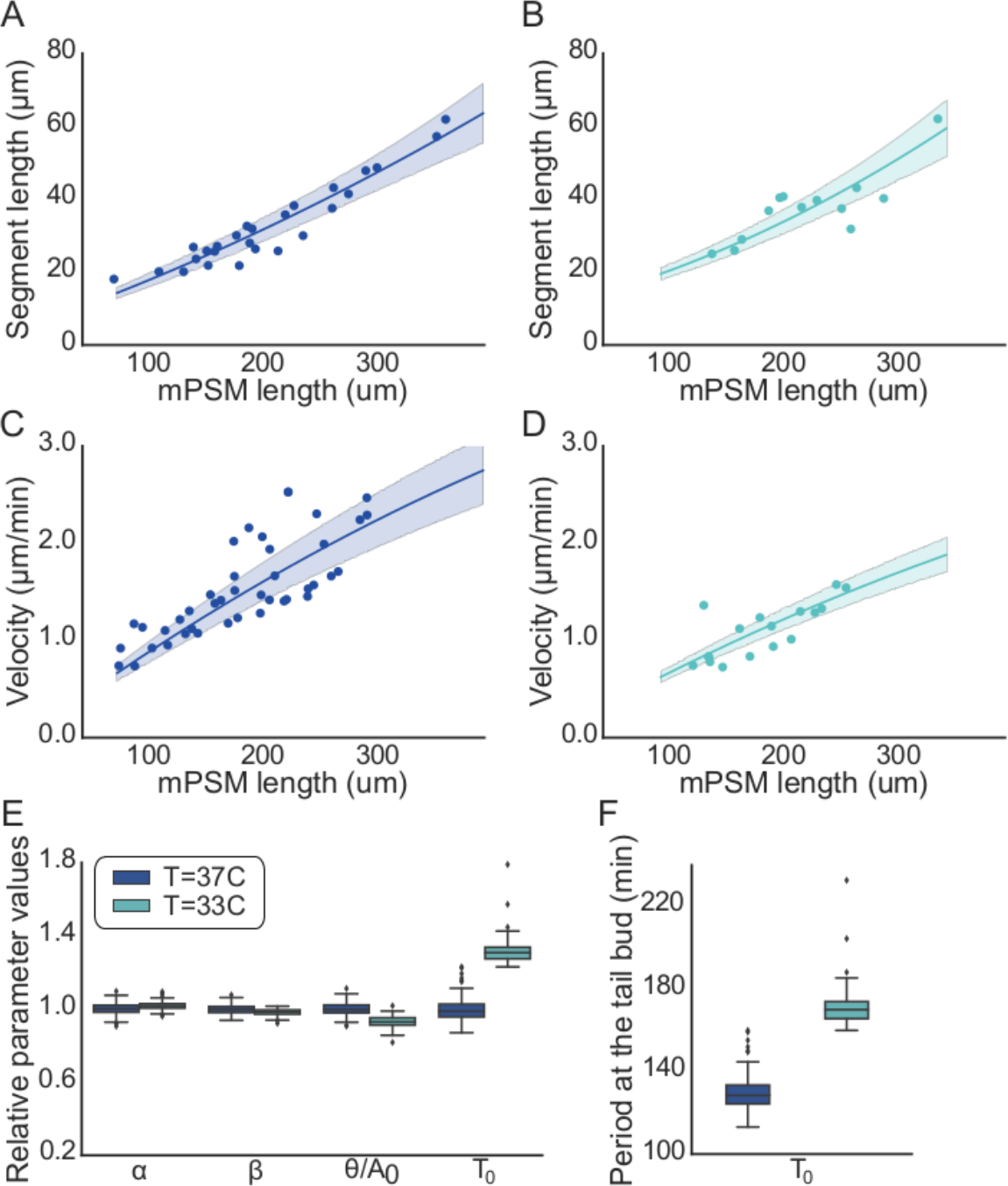
Properties of mouse segmentation in ex vivo explants at different temperatures. A) Segment size and mPSM length for explants at 37°C and B) 33°C. C) Wave velocity and PSM length for explants at 37°C and D) 33°C. The wave velocity is defined by the mPSM divided by the time to form the segment. A-D) Dots represent data from (Lauschke et al, 2013), lines represent the average fit and area represent the 95% confidence interval of the model prediction. E) Comparison of parameter values that best fit the data for explants at temperatures 37°C and 33°C. The values are normalized by the average values of explants at 37°C. Changes in the period at the center of the mPSM (T_0_) are required to fit explants from different temperatures. F) The predicted increased period for explants at lower temperatures is consistent with experimental observations of an overall period of 137.4 min and 193.4 min for explants at 37°C and 33°C (Lauschke et al, 2013). Moreover, in order to fit explants at 37°C our model requires a period at the center of the explant of around 130 min, which is consistent with the value obtained experimentally of 132 min (Lauschke et al, 2013).

Taken together, these results suggest that our framework is able to reproduce somite formation from both *in vivo* and *ex vivo* data. The critical difference between both cases is that a spatial gradient emerges in *ex vivo* explants by self-organisation, leading to a scaling of segment size with the mPSM, whereas *in vivo*, the tail bud growth controls the delayed scaling between somite size and PSM length.

## Discussion

Here, we propose a novel mechanism of spatial pattern formation that does not require any long-range interactions, such as morphogen gradients, to define the position of somite boundaries. It requires only a single cellular oscillator in each cell with a temporal modulation in the period and amplitude, as experimentally observed (Delaune et al, 2012; Shih et al, 2015; Gomez et al, 2008; Tsiaris and Aulehla, 2016). Because the oscillators have slight differences in the period and amplitude in neighboring cells, the differences in the levels between neighboring oscillators increase temporally. We propose that once this difference is large enough, the segmentation program starts. This leads to a timer mechanism, where the program of somite formation is already encrypted into the cells before they leave the tail bud region. After that, the cells only need to compare the levels of their oscillator with their neighbor’s to decide when it is time to form a new segment.

The model succeeds in reproducing the measured time-varying PSM length and somite sizes in both WT and growth disturbed mouse embryos (Fig. 3), as well as in *ex vivo* mouse explants at different temperatures (Fig. 7). Moreover, our model establishes a relationship between somite size, PSM length and growth rates (Eq. 17–20). The PSM length is proportional to the growth rate and consequently the rate of shrinkage of the PSM length is proportional to dynamic changes in the tail bud growth rate, and does not require changes in the properties of the signaling gradients, as required by the clock-and-wavefront model. According to our timer mechanism, somite size is determined at the moment the cells leave the tail bud and is proportional to the period of the oscillators at the tail bud and the tail bud growth rate at the time the cells are incorporated into the tissue (Eq. 18). In the case of constant growth rates, our model leads to the same prediction as the clock-and-wavefront, since the wavefront velocity is the same as the tail bud growth rate. However, at least in the mouse, the growth rates vary substantially during somitogenesis (Fig. 3). In the case of dynamic growth rates, because the size of the somites are defined much earlier, our model predicts a delayed scaling between PSM length and somite size, where the delay is approximately the ratio PSM/somite size (Eq. 21). We showed that experimental data from different species is in agreement with this prediction (Figure 4), supporting the idea that the somite segmentation is controlled by a timer mechanism rather than a spatial wavefront.

There are three key requirements for our mechanism to work: i) There must be an increase in period and amplitude over time. ii) There must be a link between the amplitude and period dynamics and the growth rate. iii) A molecular mechanism must exist that allows cells to sense a difference between the value of their intracellular oscillator and that of their neighbours. The first requirement, that the period and amplitude of Hes/her oscillators increase over time, has been established experimentally (Delaune et al, 2012; Shih et al, 2015; Gomez et al, 2008; Tsiaris and Aulehla, 2016), but it is not known how this increase is regulated and whether and how it may be linked to the growth rate. Experimental observations suggest a link via Wnt signalling. Thus, as cells enter the PSM, the levels of Wnt start to decrease over time due to mRNA decay (Aulehla et al, 2003). The decay of Wnt activity has been found modulate the period of Hes/her oscillations in PSM cells (Gibb et al, 2009; Wiedermann et al, 2015; Dubrulle et al, 2001; Sawada et al, 2001), and also to modulate growth at the tail bud (Amin et al, 2016). Whether Wnt also modulates the amplitude of the oscillators remains to be confirmed. These results suggest the possibility that Wnt activity link the properties of the oscillators such as the characteristic time scale of the amplitude and period gradients (α and β) with the growth rate. The coupling between the properties of the oscillators and growth has previously been shown to lead to a segmentation process that can account for PSM shrinking and growth termination (Jörg et al, 2015; Jörg et al, 2016), and in our model, it is required in order to fit data from different species (Figure 5). Moreover, such coupling suggests a possible developmental mechanism of somite size control. Species with low metabolic rates, such as snakes, have low growth rates and also slower mRNA degradation. Therefore, in these species, Wnt activity would decay more slowly and as a consequence, the changes in the properties of the oscillators would also be slower, as represented by an increased time scale of the amplitude and period gradients (α and β). As both, α and β, modulate the PSM length, but not somite size (Figure 6C), species with slower metabolism would have a larger PSM to somite size ratio and lower growth rates, as observed when comparing different species (Gomez et al, 2008; Figure 5B).

Lastly, our model requires the existence of a molecular mechanism that enables neighboring cells to compare their protein concentration to obtain their positional information. The Hes/her oscillations are part of the Notch pathway. PSM cells communicate with their neighbors via Notch/Delta signaling, and Notch signaling has been shown to control *Mesp2,* which is required to initiate somite segmentation (Takahashi et al, 2000; Morimoto et al, 2005; Yasuhiko *et al.* 2006). It is possible that as long as the differences in amplitude and period are small between oscillators, communication between neighboring cells maintains oscillations synchronized (Riedel-Kruse et al., 2007; Delaune et al., 2012; Jiang et al, 2000; Tomka et al, 2018). However, once the differences exceed a critical value, entrainment breaks. Based on this, one would expect oscillators in the PSM to remain entrained until differences become too large. The boundary of entrainment would correspond to the segmentation boundary. How cells would sense the lack of entrainment is not known. Fgf is well known to control the somite boundary (Dubrulle et al, 2001) and recent experimental evidences in zebrafish show that the spatial difference in Fgf activity is constant at the determination front (Simzek and Özbudak, 2018). In mice, Fgf signaling activity is dynamic and has been shown to be dependent on Notch activity via *Hes7* expression (Niwa et al, 2011). This suggests the possibility that Fgf and Notch work together to form a decoding mechanism of spatial differences in signaling activity between neighboring cells. The molecular mechanisms underlying such a decoder mechanism require further theoretical and experimental investigation.

## Material and methods

### Experimental data

Experimental data was obtained from previous published manuscripts. Data from *in vivo* mouse segmentation was obtained from (Tam 1981), *ex vivo* explants from (Lauschke et al, 2013) and different species from (Gomez et al, 2008). Data were extracted from original manuscripts using WebPlotDigitizer 4.1 online tool (https://apps.automeris.io/wpd/).

### Parameter values

**Table 1.** Parameter values used in the simulations, unless indicated otherwise.

| Parameters | Figure 3 (WT, MMC) | Figure 4,5,6 | Figure 7 |
| --- | --- | --- | --- |
| $\alpha$ or $\alpha'$ | 145, 100 min | 160 min | 240 $\mu\text{m}$ |
| $\beta$ or $\beta'$ | 1154, 1092 min | 1200 min | 2470 $\mu\text{m}$ |
| $g$ | from data | 1.5 $\mu\text{m}/\text{min}$ | ----- |
| $T_0$ | 81, 71 min | 100 min | 130 min |
| $\vartheta/A_0$ | 1.0, 0.92 | 1.0 | 0.05 |
| $Y$ | 37, 37 min | 30 min | ----- |

## Code availability

The simulations were evaluated in Python and all source codes are presented as Jupyter notebooks (http://jupyter.org/) for easy visualization and are freely available at: https://git.bsse.ethz.ch/iber/Publications/2018Boareto_AmplitudeModelSomitogenesis

## Parameter estimation

In order to estimate the parameters of our model that best fit the experimental data, we defined a cost function based on the Euclidean distance between the experimental and the theoretical data points. We then found the parameters that minimize this cost function by using the Python library (scipy.optimize.minimize). For more details, please see the source code: https://git.bsse.ethz.ch/iber/Publications/2018Boareto_AmplitudeModelSomitogenesis

## Acknowledgements

The authors thank members of Iber lab for comments. M.B. thanks T. A. Amor for helping with the figures. This work was supported by the SystemX.ch of the Schweizerischer Nationalfonds zur Förderung der Wissenschaftlichen Forschung (Swiss National Science Foundation) and the NeuroStemX project.

## Author contributions

Conceptualization: M.B., T.T. and D.I.; Performed research: M.B. and T.T.; Developed the theoretical framework: M.B.; Writing: M.B. and D.I.; Funding acquisition and project administration: D.I.

## Supplementary Information

**Figure S1.**
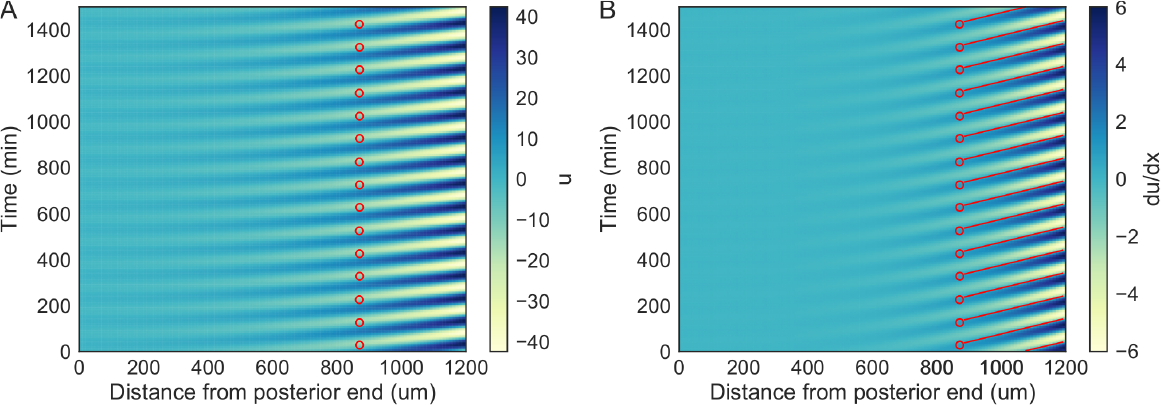
Properties of the oscillators during segmentation. A) Levels of the oscillators and B) its spatial derivative as a function of time and position related to the posterior end. Red dots and lines represent the position of formation of a new segment, i.e., where du/dx = θ. Note that the caudal cells reach the threshold before the rostral cells within a somite length. The distance between the previous somite and the first most caudal cell to reach the threshold define the somite size.

**Figure S2.**
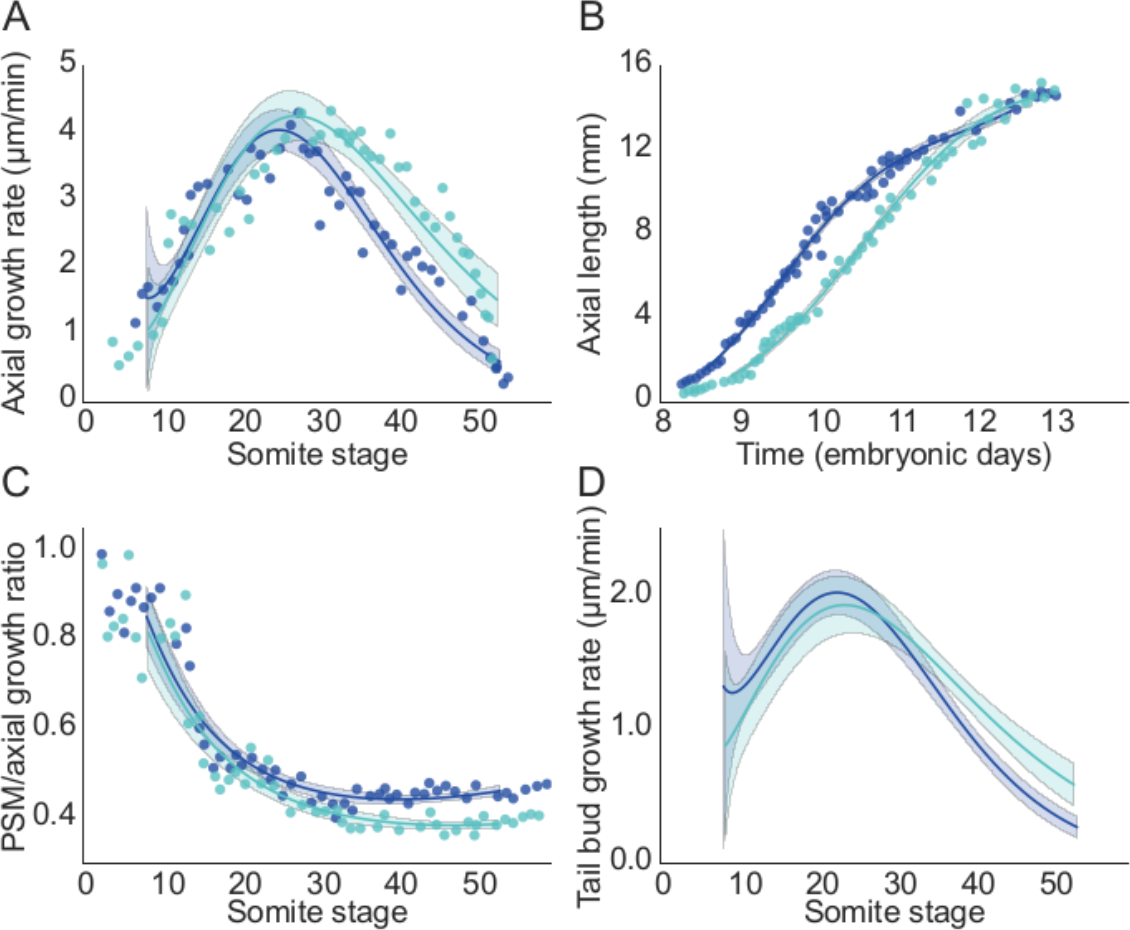
Growth profiles of WT and MMC-treated embryos. A) Axial growth rate for different somite stages. B) Axial length for different embryonic times. C) Fraction of growth rate due to tail bud elongation for different somite stages. D) Tail bud growth rate for different somite stages inferred by scaling the axial growth rate with the relative growth due to tail bud elongation. A-C) Points represent data from (Tam 1981), solid line represents average fit and area represents 95% confidence interval of a bootstrap fit. Dark blue represents WT and light green represents MMC-treated embryos.

**Figure S3.**
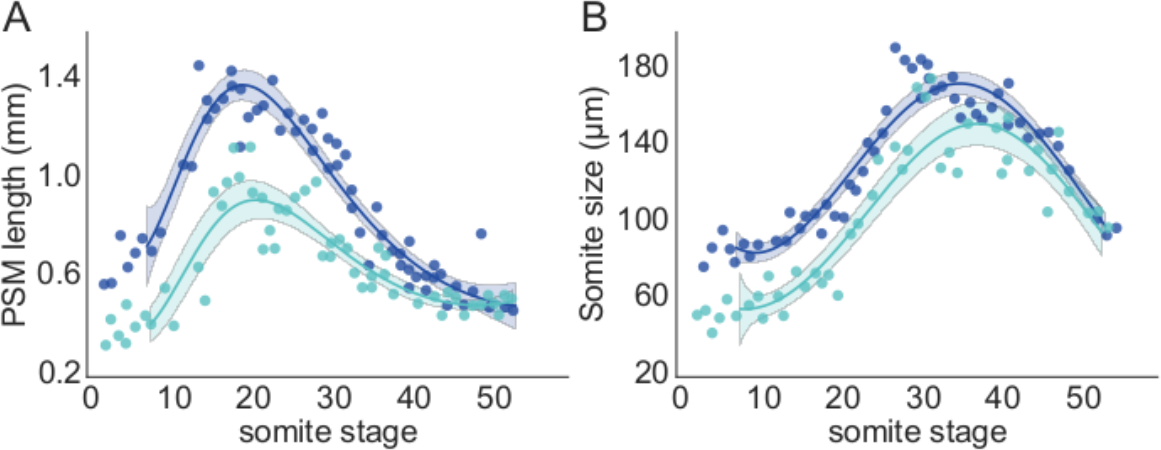
PSM length and somite size profile of WT and MMC-treated embryos. A) PSM length for different somite stages. B) Somite size for different somite stages. Points represent data from Tam 1981, solid line represents average fit and area represents 95% confidence interval of a bootstrap fit. Dark blue represents WT and light green represents MMC-treated embryos.

**Figure S4.**
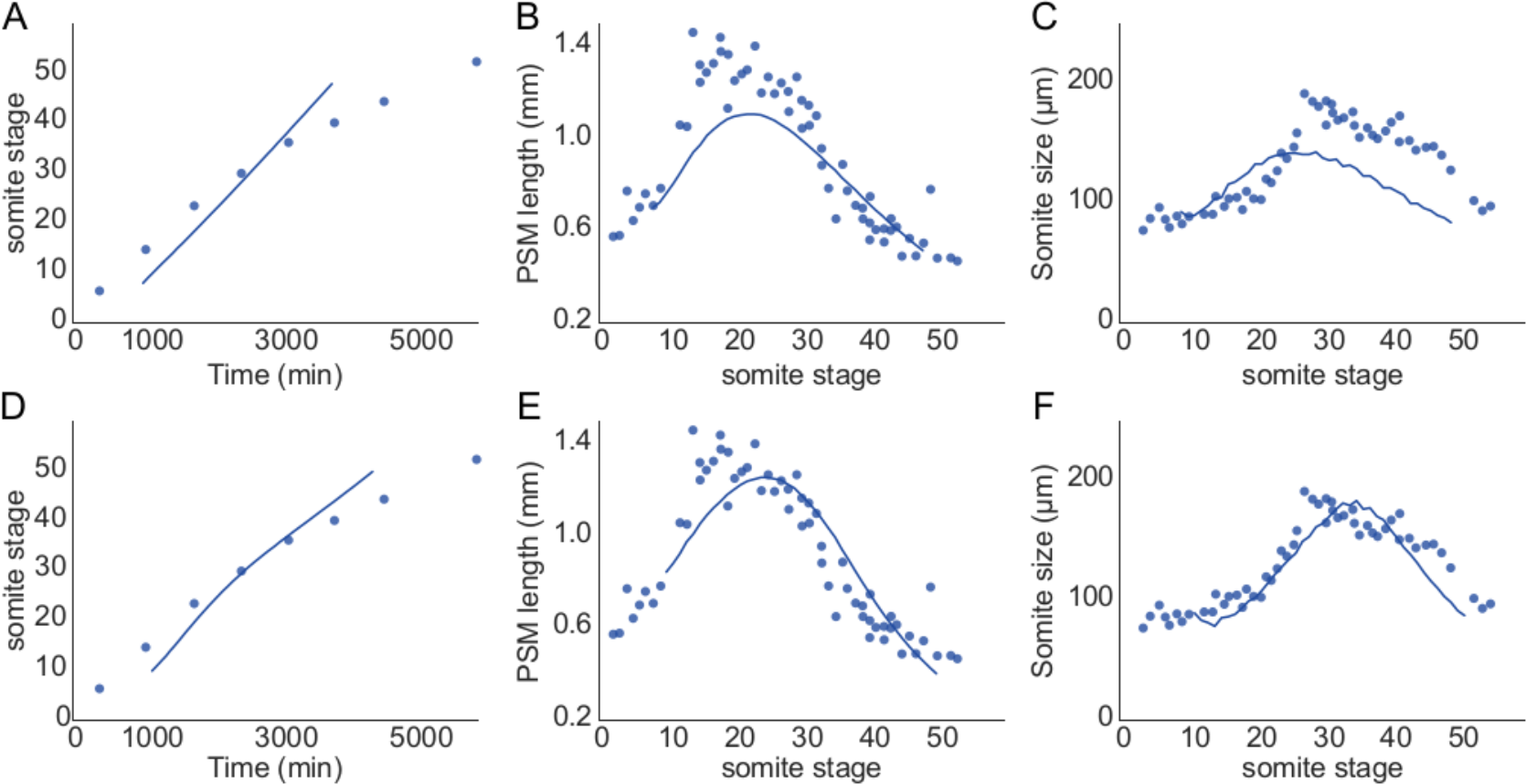
Fit of mouse in vivo segmentation data. A,D) Relationship between somite stage and embryonic time. B-E) PSM length at different somite stages. C,F) Somite size at different somite stages. Circles represent data from (Tam 1981) and lines represent the fit from the modeling using the average value of tail bud growth rate as input. A-C) Modeling results when considering constant period and amplitude at the tail bud. D-F) Modeling results when considering that the period and amplitude at the tail bud are dynamic and dependent on the tail bud growth rate.

**Figure S5.**
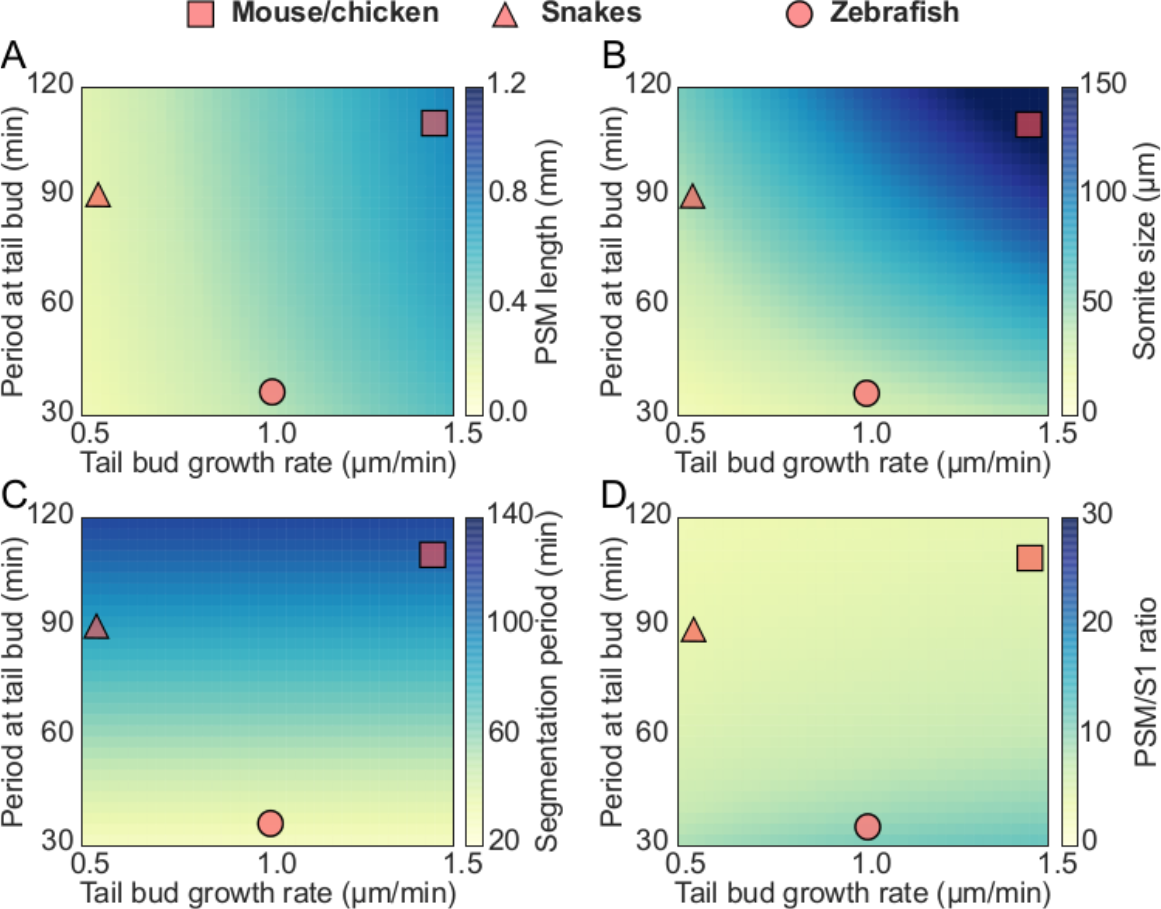
A) PSM length, B) somite size, C) segmentation period and D) PSM/somite size ratio for different values of growth rate and period at the tail bud. The characteristic length of amplitude and period gradient is remained constant (same values of Figure 3, see Table 1). Note that in this case, our model would predict that snakes would have a much smaller PSM length (around 300 μm). This is in contrast with experimental evidences showing that snakes have a PSM length of around 1200 μm (Gomez et al, 2008).

**Figure S6.**
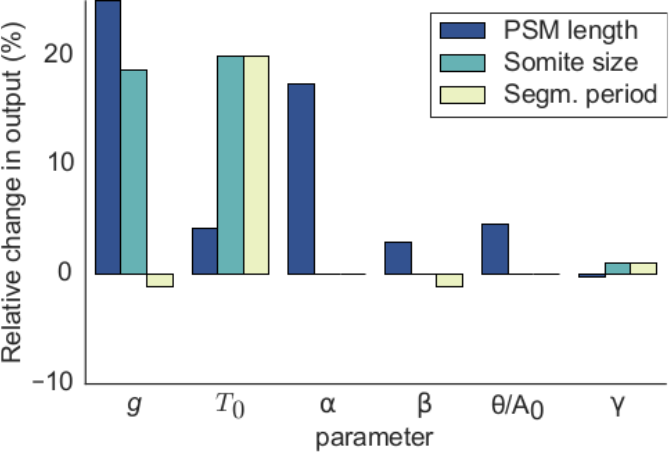
Sensitive analysis by considering the period and amplitude at the tail bud is dependent on the growth rate. The extra parameter γ has little effect on PSM length, somite size and segmentation period.

## Static period gradient in *ex vivo* explants

In our analysis, we represented the period gradient observed in *ex vivo* explants as static, i.e., the period gradient does not change temporally, only spatially. This is in contrast to what is proposed in (Lauschke et al, 2013), where the authors infer a dynamic, exponentially increasing period gradient. We reanalyzed the data from (Lauschke et al, 2013) and concluded that their data is more consistent with a static period gradient. For this reason, we consider the static case in our analysis.

In our model, we consider a spatial period gradient that is constant in time. In case of such a static period gradient, the phase (*ϕ*) and the slope of the phase-gradient 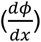 are linear in time and are given by: *ϕ* = *ωt* and 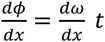, respectively, where *ω* = *ω*(*x*) represents the frequency of oscillations. To test whether the period is static or dynamic, one can therefore plot the phase-gradient against time and evaluate the slope. If the slope is constant in time, then the spatial frequency gradient, 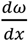, is constant in time, which would indicate a static period and *vice versa*. Lauschke and co-workers observed a linear relationship between the number of oscillations and time (Figure S7), and the phase-gradient can therefore be plotted against the oscillation number.

**Figure S7.**
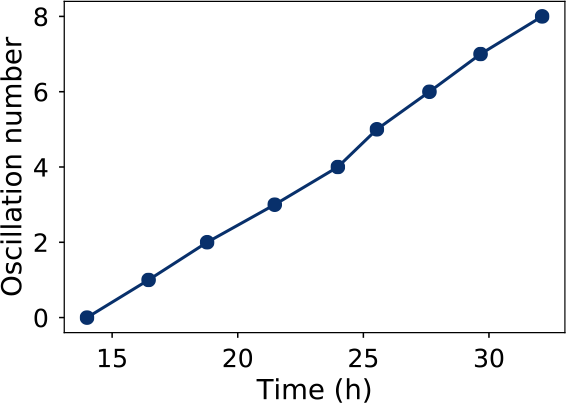
Linear relationship between oscillation number and time. Blue dots represent the time of formation of a new segment (digitalized from Figure 4a, Lauschke et al, 2013). The slope of the curve represents the time it takes to form a new segment and is approximately the period at the center of the periphery (132 min, Lauschke et al, 2013, Figure Supplementary 4).

Figure S8A shows the plot of the phase-gradient against the oscillation number. Here, the phase-gradient is plotted on a logarithmic scale. At first sight, the data may suggest a linear relationship on the logarithmic scale, and thus a non-linear relationship between the phase-gradient and time and a time-dependent period. However, a linear relationship (red curve) fits the data essentially as well, in particular when evaluated on a linear scale (Figure S8B). Why would a linear and an exponential curve be so similar? A linear relationship is clearly different from an exponential relationship only if the exponent in the exponential function is sufficiently large and the time period over which the function is observed is sufficiently long. This is not the case for the reported segmentation data: the experimentally determined time-dependency of the phase gradient can be fitted just as well by a linear function (Figure S8, red line), and we confirm that our model recapitulates this relationship just as well (Figure S9, red).

We note that the Lauschke and co-workers previously showed that the period at the center remains constant during the segmentation process (Lauschke et al, 2013, Figure Supplementary 4). We would argue that in light of this experimental observation, it is rather likely that the same applies also to the rest of the domain such that a static frequency gradient would be a better representation of the data.

**Figure S8.**
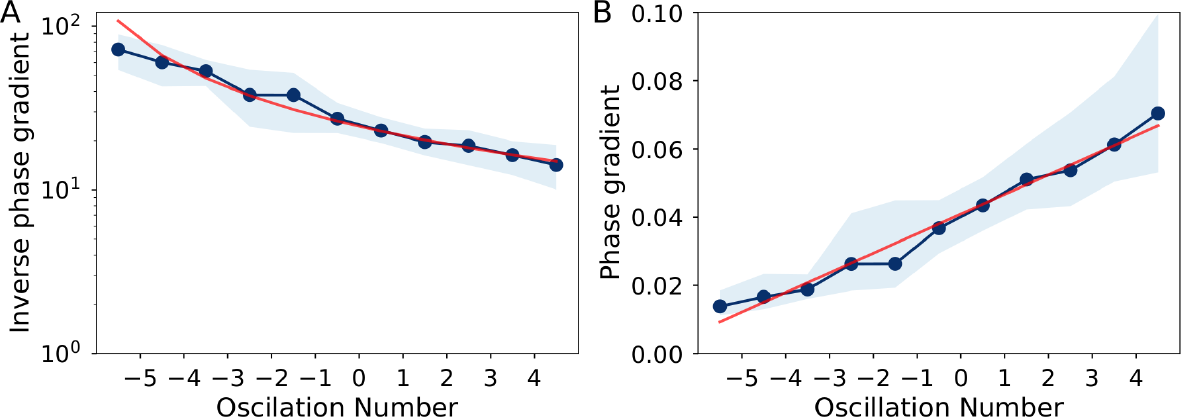
Linear fit of spatial phase-gradient. A) Inverse of the spatial phase-gradient as a function of oscillations number, as represented in (Lauschke et al, 2013, Figure 4b). Note that oscillation number is linearly proportional to time (Figure S7). B) Spatial phase-gradient as a function of oscillations number. Blue line represents experimental data and area represents the experimental standard deviation and red line represents a linear fit. Negative oscillation numbers represent a spatial projection of incomplete segmentations (Lauschke et al, 2013).

**Figure S9.**
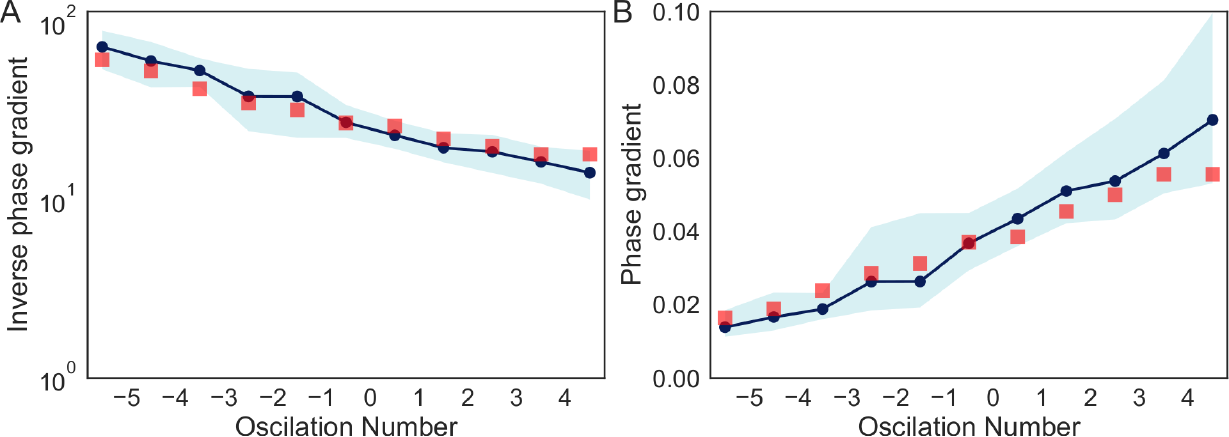
Spatial phase-gradient in time. A) Inverse of the spatial phase-gradient as a function of oscillations number, as represented in (Lauschke et al, 2013, Figure 4b). B) Spatial phase-gradient as a function of oscillations number. Blue line represents experimental data and area represents the experimental standard deviation. Red dots represent the relationship obtained by our model.

